# GABA_B_- GluK1 kainate receptor interplay modulates amygdala excitability and behavioral response to chronic stress

**DOI:** 10.1101/2024.01.15.575620

**Authors:** Maria Ryazantseva, Maj Liiwand, Sari E. Lauri

## Abstract

Amygdala hyperexcitability is a hallmark for stress-induced anxiety disorders. Stress-associated changes in both principal neurons and interneurons contribute to the increased excitability, but how exactly these mechanisms perturb function of behaviorally relevant circuits in the amygdala remains unclear. Here, we show that GluK1 subunit-containing kainate receptors in parvalbumin (PV) interneurons maintain high GABA release and control excitability of lateral amygdala (LA) principal neurons via tonic GABA_B_-receptor-mediated inhibition. Downregulation of GluK1 expression in PV interneurons after chronic restraint stress (CRS) releases the tonic inhibition and increases excitability of LA principal neurons. Stress-induced LA hyperexcitability facilitates glutamatergic transmission selectively to central amygdala PKCδ-expressing neurons, implicated in fear generalization. Consistent with significance in anxiogenesis, absence of GluK1- GABA_B_ regulation confers resilience against CRS-induced LA hyperexcitability and anxiety-like behavior. Our data reveal a unique novel mechanism involving an interplay between glutamatergic and GABAergic systems in the regulation of amygdala excitability in response to chronic stress.

## Introduction

Chronic stress produces lasting structural and functional changes in various areas of the brain, which contribute to the neuropathology of stress-related psychiatric diseases. Amygdala is one of the key structures implicated in the stress response in both humans and animal models. Stress hormones increase the discharge rates and firing synchrony of basolateral amygdala (BLA) neurons, which facilitates fear and aversive learning that is critical for animal survival (Pare and Headley, 2023). During prolonged and severe stress, however, these mechanisms may become maladaptive and result in sustained hyperexcitability of BLA principal neurons (Roozendaal et al., 2009). Excessive activity of the BLA disturbs regulation of downstream brain areas involved in emotional responses and associates with various behavioral disorders characterized by unwarranted fear and anxiety (reviewed by Tovote et al., 2015; Calhoon and Tye 2015; Prager et al., 2017; Sharp et al., 2017; Zhang et al., 2021).

The cellular mechanisms underpinning stress-induced amygdala hyperexcitability have been widely studied and shown to involve alterations in both, the intrinsic properties of the BLA principal neurons and the surrounding network (reviewed by Zhang et al., 2021). In particular, chronic stress perturbs the function of parvalbumin–expressing (PV) GABAergic interneurons, which mediate perisomatic inhibition of the principal neurons and tightly control their excitability and local network dynamics (Hu et al., 2014; Prager et al., 2017; Babaev et al., 2018). PV interneurons respond to stress in an age- and sex-specific manner (Woodward and Coutellier, 2021) and their activity in BLA has been directly linked to anxiety-like behaviors in mice (Luo et al., 2020). Yet, how exactly stress-dependent changes in PV interneurons contribute to the hyperexcitability of the principal neurons and how this affects the intra-amygdaloid circuit regulating fear-related behaviors remains poorly understood.

To get further insight into the mechanisms underlying aberrant behaviors induced by chronic stress, we have here focused on the trisynaptic circuit involving PV interneurons and principal neurons (PN) in the lateral amygdala (LA), as well as their GABAergic PKCδ−expressing target neurons in the centrolateral (CeL) amygdala. CeL PKCδ neurons are in a key position in controlling amygdala output and have been strongly implicated in regulation of anxiety and fear generalization (Tye et al., 2011; Cai et al., 2014; Botta et al., 2015). We show that the excitability of LA PN’s is regulated by tonic GABA_B_-mediated inhibition that is maintained by kainate-type glutamate receptors (KARs), facilitating asynchronous GABA release in PV interneurons. This mechanism is lost after chronic stress, due to downregulation of GluK1 KAR subunit expression specifically in PV interneurons. The increase in BLA excitability in the absence of tonic GluK1-GABA_B_-mediated inhibition affected glutamatergic synaptic transmission to CeL in a cell-type specific manner, shifting the balance of excitatory drive towards PKCδ neurons. Consistent with significance in fear generalization and anxiogenesis, ablation of GluK1 specifically in PV interneurons GluK1 expression conferred resilience against CRS-induced LA hyperexcitability and anxiety-like behavior.

## Results

### Loss of GABA_B_-receptor-mediated tonic inhibition contributes to amygdala hyperexcitability after chronic stress

To study the effect of chronic stress on amygdala circuitry and anxiety-like behaviors, we submitted mice to chronic restraint stress protocol (CRS) involving 1 h immobilization in a ventilated tube during ten consecutive days. As expected, CRS associated with significant loss of weight (Figure 1A) as well as increase in the anxiety-like behavior in the open field test. Thus, the CRS-exposed mice spent significantly less time in the center area of the open field arena as compared to controls (Figure 1B). In addition, CRS-exposed mice showed higher locomotory activity, indicated by the total distance traveled during the test (Figure 1B). Yet, consistent with an anxiety-like phenotype, the ratio of distance traveled in the center field to the total distance in CRS group was lower as compared to controls (48 ± 3 % and 62 ± 4 %, respectively, p<0.05, unpaired t-test).

**Figure 1.**
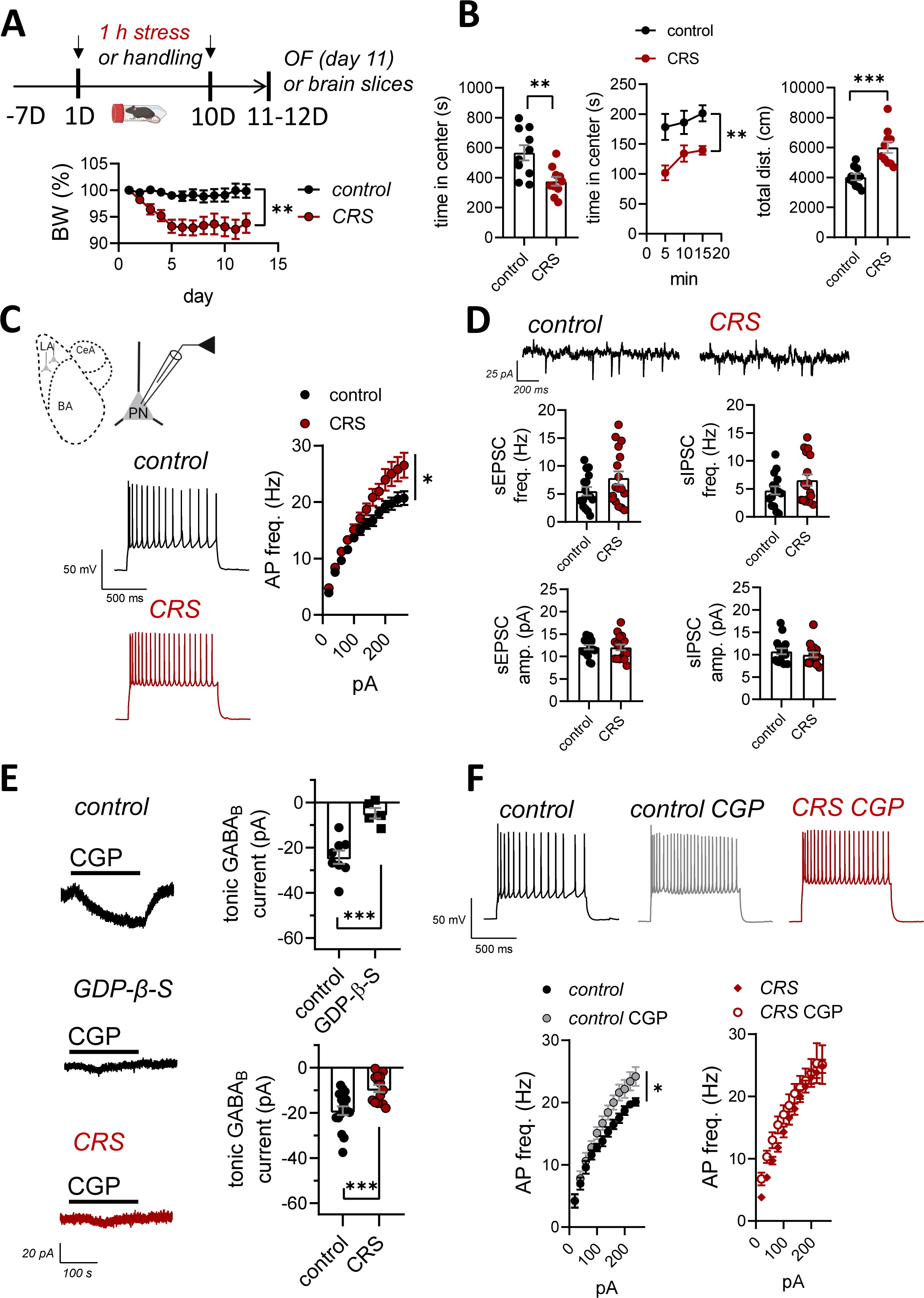
Loss of GABA_B_ R-mediated tonic inhibition contributes to amygdala hyperexcitability after chronic stress. (A) A scheme of the experimental protocol (top) and a graph illustrating body weight changes during the chronic restraint stress (CRS, n=10, control n=10; RM ANOVA F_(1, 18)_ = 10.54, **p=0.0045) (B) Results of the open field (OF) test. The graphs show the total time spent in the center area of the open field (OF) and the center time in 5 min bins (control, n=10, CRS, n=10, t-test t=3.232, df=18, **p=0.0046; RM ANOVA F_(1, 18)_=10.45, **P=0.0046). Total distance traveled during OF test (t-test, t=4.546, df=18, ***p=0.0003). (C) Action potential (AP) firing rate of principal neurons (PN) in the lateral amygdala (LA) in response to depolarizing current steps, recorded from brain slices of control and CRS-exposed mice (control, n=17 (8 mice), CRS, n=19 (7 mice); RM ANOVA F _(1, 29)_ = 4.877, *p=0.035). The example traces illustrate the response to 240 pA step current. (D) Example traces for recordings of spontaneous synaptic activity from LA PN neurons, using a low- chloride containing electrode filling solution at -50 mV holding potential. Under these conditions, sEPSCs and sIPSCs are observed as inward and outward currents, respectively. Pooled data on the sEPSC and sIPSC frequency and amplitude for control and CRS groups (control, n=16 (8 mice), CRS, n=17 (7 mice); sEPSC: t-test t=1.296, df=31, p= 0.205; sIPSC: t-test t=1.476, df=31, p= 0.15). (E) Tonic GABA_B_ receptor-mediated currents recorded from LA PNs in response to the application of GABA_B_ antagonist CGP55845 (10 μM), in control and CRS-exposed mice as well as in the presence of GDP-β-S in a control mouse. All recordings were done in the presence of 50 μM of D-AP5, 200 μM picrotoxin, and 50 μM GYKI 53655 to block NMDA, GABA_A,_ and AMPA receptors, respectively. Pooled data on the maximal amplitude of the GABA_B_ current under control conditions and in the presence of GDP-β-S (750 µM) in the electrode filling solution (control, n=8 (3 mice), GDP-β-S n=5 (3 mice); t-test, t=4.599, df=11, ***p=0.0008). Amplitudes of the tonic GABA_B_ current in control and CRS- exposed animals (control, n=17 (7 mice), CRS, n=14 (3 mice); t-test t=3.786, df=29, ***p=0.0007). (F) Effect of GABA_B_ antagonism on firing frequency of LA PNs in control and CRS-exposed mice. Action potential frequencies in response to depolarizing current steps were recorded from brain slices of control and CRS-treated animals, at control conditions and in the presence of CGP55845 (5 µM) (control, n=10 (4 mice), control+CGP55845, n=10 (4 mice); RM ANOVA, F_(2, 26)_=3.498, *p=0.045; CRS, n=17 (6 mice), CRS+CGP55845, n=9 (3 mice); RM ANOVA, F_(1, 24)_ = 1.824, p=0.1894). Example traces show the response to 240 pA current step. All the data are presented as mean ± SEM

After validating the CRS protocol, we went on to study the excitability of LA principal neurons (PNs) using whole-cell current-clamp recordings in acute brain slices from control and CRS-exposed mice. Consistent with previous reports (Rosenkranz et al., 2010; Zhang et al., 2019), the firing rate of the LA PN’s in response to depolarizing current steps was significantly higher in CRS-exposed mice as compared to controls (Figure 1C). This effect was associated with lower amplitude of medium duration after-hyperpolarizing current (AHP) (Rosenkranz et al., 2010; Rau et al., 2015; Zhang et al., 2019) and more depolarized resting membrane potential (control 61.44 ± 1.47; CRS 57.10 ± 1.32 mV, p=0.03, t-test) (Supplementary Figure 1A,B). No other differences in the properties of action potentials were detected between the groups (AP half-width, AP threshold, amplitude of the fast AHP current; Supplementary Figure 1A).

Previously, it has been reported that CRS results in significant alterations in both, glutamatergic and GABAergic synaptic inputs to the BLA principal neurons (e.g. Wei et al., 2018; Luo et al., 2020; Liu et al., 2020; Zhang et al., 2021). However, we observed no significant differences between the control and CRS groups in the frequency or amplitude of spontaneous glutamatergic or GABAergic synaptic responses (sEPSCs and sIPSCs, respectively), that were recorded simultaneously under voltage clamp from LA PNs (Figure 1D). Since GABAergic input originates from distinct subtypes of interneurons that might be differentially regulated by CRS, we went on to record sIPSCs under modified conditions to be able to distinguish fast (perisomatic) and slower (distal) synaptic events based on their kinetics. Despite of the increased resolution, we observed no differences between control and CRS groups in the frequency, amplitude, rise time, or decay time distribution of sIPSCs, recorded from LA principal neurons using high-Cl containing electrode filling solution in the presence of glutamatergic antagonists (Supplementary Figure 1 C,D).

GABAergic activity can influence the target cells also via extrasynaptic, tonic inhibition (Farrant and Nusser, 2005). Chronic stress reduces tonic inhibition mediated by extrasynaptic GABA_A_-receptors in the BLA, which contributes to hyperexcitability of BLA PN’s (Liu et al., 2014; Qin et al., 2022). Recently, also GABA_B_-receptors have been implicated in tonic inhibition of the BLA principal neurons (Mackay et al., 2019; Marron Fernandez de Velasco et al., 2023), yet their role in stress-induced amygdala hyperexcitability is not known. We observed that application of CGP55845, a selective GABA_B_-receptor antagonist, resulted in a significant shift in the holding current of the LA principal neurons, and this was fully blocked when G-protein coupled signaling was prevented by inclusion of the GDP-β−S in the recording electrode (Figure 1E). This CGP55845 sensitive tonic GABA_B_ current was significantly smaller in the CRS-exposed mice as compared to controls (Figure 1E). Furthermore, current clamp recordings indicated that application of CGP55845 resulted in a significant increase in the firing rate of the LA PNs in response to depolarizing current steps in control slices, but not in slices from CRS-treated mice (Figure 1F). Together, these data demonstrate that loss of tonic GABA_B_ receptor-mediated inhibition contributes to the increased firing rate of LA PNs after chronic stress.

### GABA_B_R-dependent tonic inhibition of LA PNs is driven by GluK1 kainate receptors in PV interneurons

GluK1 subunit containing kainate receptors (KARs) have been previously shown to regulate both phasic and tonic GABA_A_ receptor-mediated inhibition in the BLA (Braga et al., 2003; Wu et al., 2007; Englund et al., 2021), yet the subtypes of GABAergic interneurons responsible for this regulation are not known. Recently, we found that GluK1 KARs are endogenously active in PV interneurons in the LA and contribute to their high excitability (Englund et al., 2021). To investigate the role of these KARs in regulation of tonic inhibition of LA PNs, we performed a set of pharmacological experiments in mice with floxed GluK1 gene (*Grik1*^fl/fl^, controls) and mice lacking GluK1 expression selectively in PV interneurons (PV-Cre::*Grik1^fl^*^/fl^).

Tonic GABA_A_ currents, recorded in response to application of bicuculline (25 µM) in LA PNs, were not significantly different between the genotypes (Figure 2A). Interestingly, however, the CGP55845 sensitive tonic GABA_B_ receptor-mediated current was smaller in mice lacking GluK1 expression selectively in PV interneurons as compared to controls (Figure 2B). Furthermore, while in control mice, the tonic GABA_B_ receptor-mediated current was substantially reduced in response to application of ACET (200 nM, Figure 2B), a selective antagonist of GluK1 KARs (Dargan et al., 2009), ACET had no effect on CGP55845 induced currents in PV-Cre::*Grik1^fl^*^/*fl*^ mice (Figure 2B). These differences were not due to the loss of postsynaptic GABA_B_ receptors in the mutant mice, as the currents induced by the GABA_B_ agonist SKF97541 in LA PNs were similar between the genotypes (Supplementary Figure 1E).

**Figure 2.**
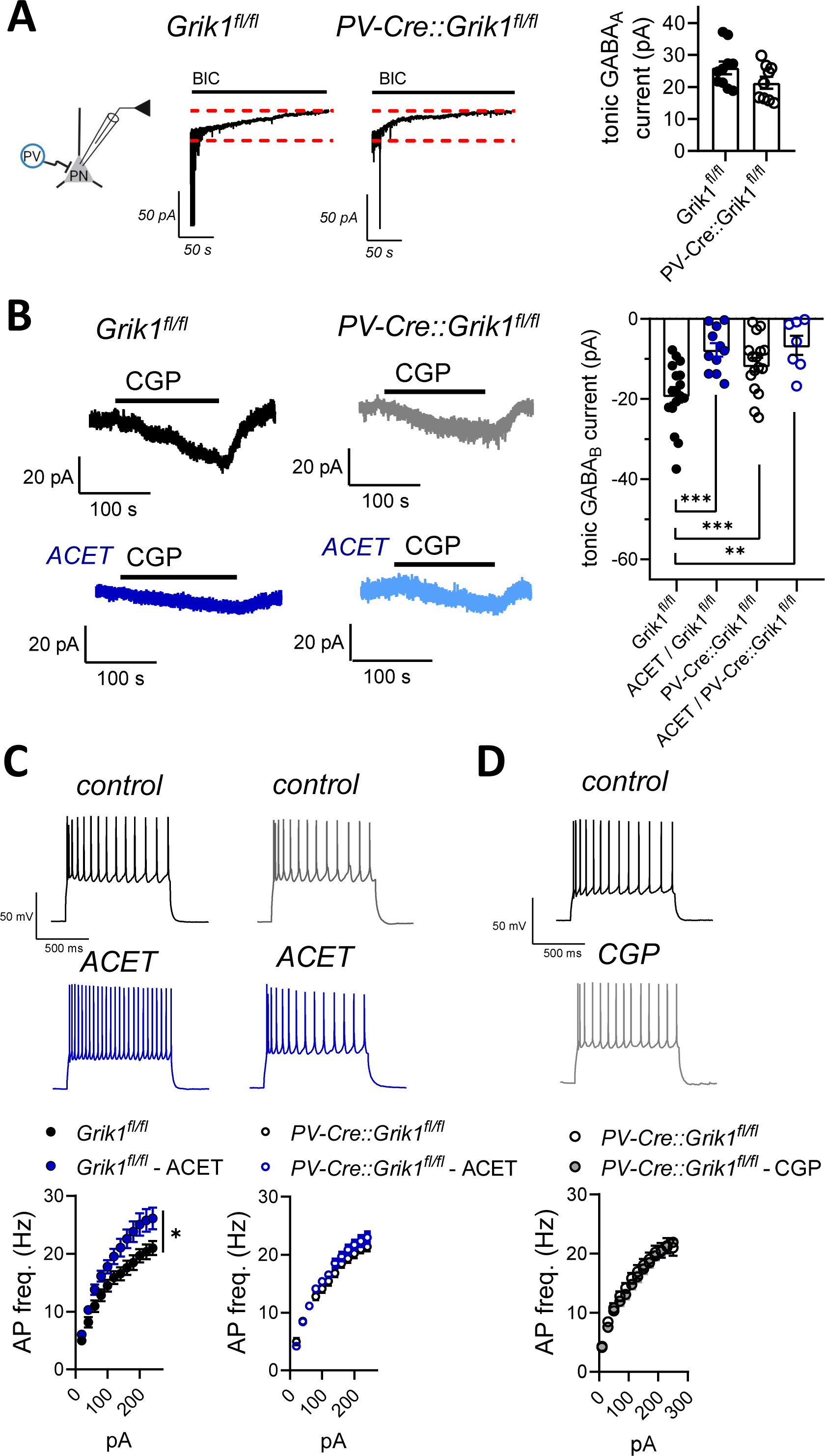
GABA_B_ R-dependent tonic inhibition is driven by GluK1 kainate receptors in PV interneurons. (A) Tonic GABA_A_ currents recorded as a change in the holding current in response to the application of 25 μM bicuculline in LA PNs, in slices from mice with floxed GluK1 gene (*Grik1*^fl/fl^, control) and mice lacking GluK1 expression selectively in PV interneurons (PV-Cre::*Grik1*^fl/fl^). All recordings were done at -90 mV holding potential in the presence of 50 μM of D-AP5, and 50 μM GYKI 53655 to block NMDA, and AMPA receptors, respectively. The graph illustrates averaged data on the amplitude of the tonic GABA_A_ current (*Grik1*^fl/fl^: n=10 (4 mice); PV-Cre::*Grik1*^fl/fl^: n=9 (3 mice); t-test, t=1.681, df=17, p= 0.11). (B) Tonic GABA_B_ currents recorded from LA PNs in slices from mice with floxed GluK1 gene (*Grik1*^fl/fl^, control) and mice lacking GluK1 expression selectively in PV interneurons (PV-Cre::*Grik1^fl^*^/fl^), before and after ACET application (200 nM). All recordings were done at -50 mV holding potential in the presence of 50 μM of D-AP5, 200 μM picrotoxin, and 50 μM GYKI 53655 to block NMDA, GABA_A,_ and AMPA receptors, respectively. The amplitude of the tonic current was measured as a change in the holding current in response to the application of 10 μM CGP55845. The graph illustrates averaged data on the amplitude of the tonic GABA_B_ currents under various conditions (Grik1^fl/fl^: control, n=17 (5 mice), ACET, n=11 (4 mice); PV-Cre::Grik1^fl/fl^: control, n=16 (4 mice), ACET, n=7 (3 mice); Dunnett test, ***p<0.001, **p=0.0095). (C) Action potential firing rate of LA PNs in response to depolarizing current steps under control conditions and in the presence of ACET (200 nM). Recordings were done in brain slices from *Grik1*^fl/fl^ and PV-Cre::*Grik1*^fl/fl^ mice (*Grik1*^fl/fl^: control, n=17 (5 mice), ACET, n=17 (5 mice), RM ANOVA, F_(1, 32)_ = 5.967, *p=0.02*; PV-Cre::Grik1*^fl/fl^, control, n=24 (8 mice), ACET, n=22 RM ANOVA, F_(1, 44)_ = 1.534, p= 0.22). (D) Action potential (AP) firing rate of LA PNs in response to depolarizing current steps under control conditions and in the presence of CGP55845 (5µM), in PV-Cre::*Grik1*^fl/fl^ mice (control, n=15 (3 mice), CGP55845, n=13 (3 mice), RM ANOVA, F_(1, 44)_ = 0.1779, p=0.6766). The example traces in C and D illustrate the response to the 240 pA current step. All the data is presented as mean ± SEM. See Supplementary data 2 for the membrane properties of LA principal neurons in *Grik1*^fl/fl^ and PV-Cre::*Grik1*^fl/fl^ mice.

Consistent with GluK1 KARs contributing to tonic inhibition of LA PNs, GluK1 antagonism by ACET resulted in a significant increase in the firing rate of the LA PNs in response to depolarizing current steps in control mice (Figure 2C). However, ACET had no effect when the same experiment was repeated in mutant mice lacking GluK1 expression in the PV interneurons (Figure 2C). Furthermore, in contrast to controls (Figure 1 F), GABA_B_ antagonism with CGP55845 had no effect on the excitability of LA PN’s in PV-Cre::*Grik1^fl^*^/fl^ mice (Figure 2D). Consistent with loss of tonic GABA_B_-mediated inhibition, the resting membrane potential of the LA PN’s in PV-Cre::*Grik1^fl^*^/fl^ mice was slightly higher as compared to controls (-57.46±1.8 mV vs -61.67±0.97 mV, respectively, p=0.047, unpaired t-test; Supplementary data 2). However, we detected no differences between the genotypes in LA PN excitability, possibly due to developmental compensation (Figure 2C; RM ANOVA for *Grik1^fl^*^/fl^ vs PV-Cre::*Grik1^fl^*^/fl^, F _(1, 36)_ = 0.6403, p= 0.429). Together, these data indicate that GluK1 KARs, located in the PV interneurons, are physiologically activated to regulate excitability of the LA PNs via GABA_B_ receptor-mediated tonic inhibition.

### GluK1 KARs facilitate asynchronous GABA release from PV interneurons

KARs have been implicated in regulation of GABA release (e.g. Rodriquez-Moreno et al., 1997; Jiang et al., 2001; Cossart et al., 2001; Daw et al., 2010; Lourenco et al., 2010; Wyeth et al., 2017), yet no direct evidence on GluK1 KARs regulating release in PV interneurons exists. Therefore, we went on to investigate whether KARs regulate action potential-dependent and/or asynchronous GABA release from PV interneurons and thereby contribute to the ambient levels of GABA, mediating tonic inhibition. To this end, we activated PV interneurons in the BLA by using cell-type specific optogenetic stimulation and recorded pharmacologically isolated GABAergic responses from LA principal neurons (Figure 3A). GluK1 KAR antagonism with ACET (200 nM) significantly reduced the amplitude of light-evoked IPSC and increased paired-pulse facilitation (Figure 3B), consistent with presynaptic GluK1 KARs facilitating GABA release in PV interneurons in the LA.

**Figure 3.**
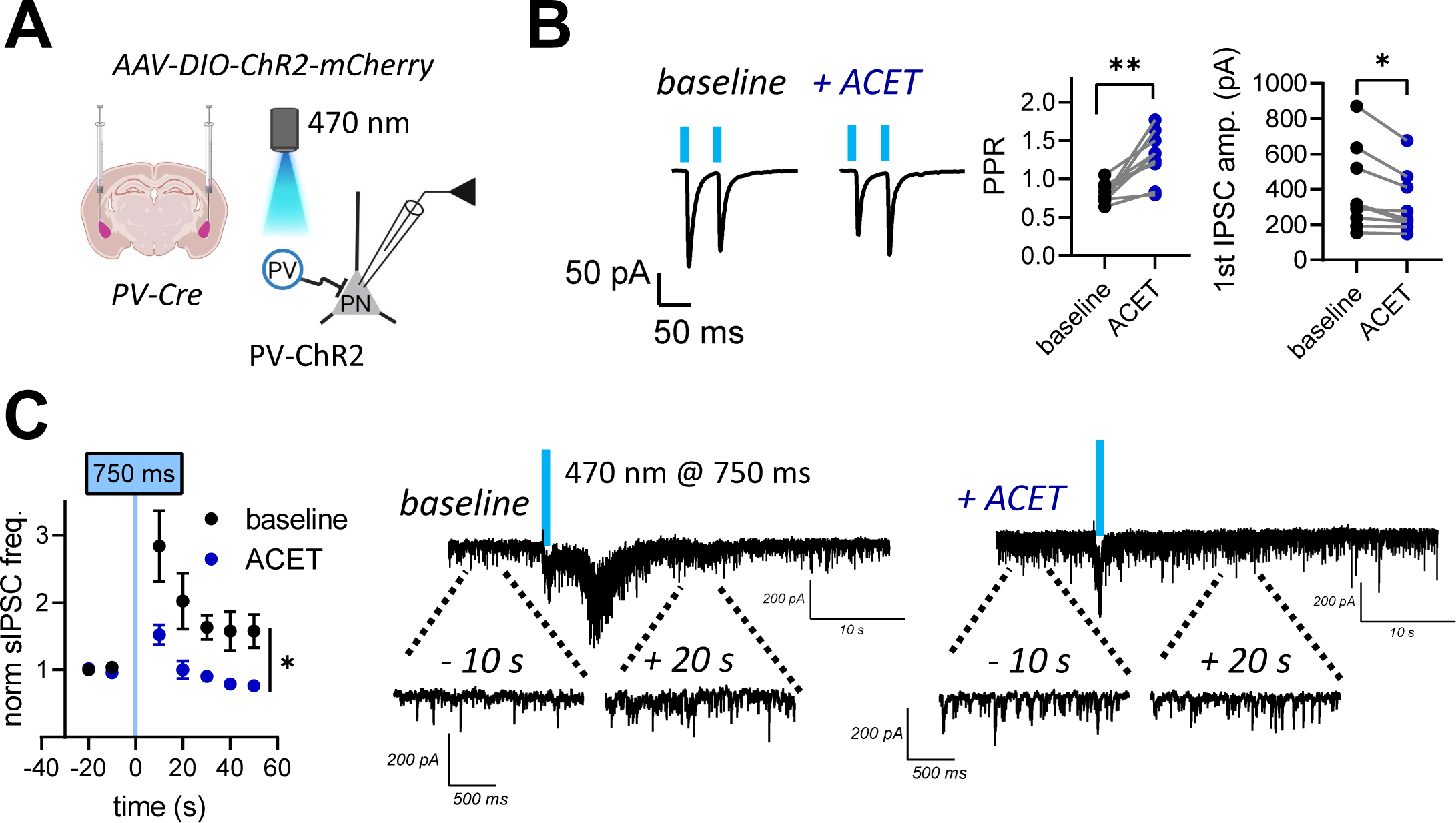
GluK1 kainate receptors regulate action potential-induced and asynchronous GABA release in PV interneurons. (A) A scheme of the experimental protocol (left). AAV viral vectors encoding for Cre-dependent ChR2 were injected to BLA of PV-Cre mice for PV neuron-specific expression of ChR2. GABAergic responses were recorded from LA PNs in response to light stimulation. (B) Examples of IPSCs, evoked by paired-pulse stimulation with 470 nm blue light before and after 200 nM ACET application. Pooled data on the effect of ACET on the 1st IPSC amplitude and paired-pulse ratio (PPR) (n=9 (5 mice), PPR, t=4.567, df=8, **p= 0.0018; IPSC, t=3.246, df=8, *p=0.0118 paired t- test). (C) Asynchronous barrage of IPSCs recorded from LA PNs in response to 750 ms opto-stimulation of PV interneurons, before and after application of 200 nM of ACET. Pooled data illustrating the frequency of IPSCs, normalized to the level before opto-stimulation and analysed in 10 min bins under control conditions and in the presence of ACET (n=8 (6 animals), RM ANOVA, F _(1, 7)_ = 11.49, *p= 0.012). All the data presented as mean ± SEM

Asynchronous GABA release from PV interneurons was evoked by prolonged light-induced depolarization, which induced a sustained increase in the frequency of sIPSCs in LA principal neurons (Figure 3C). As shown previously (Englund et al., 2021), application of ACET resulted in a significant increase in the basal sIPSC frequency in LA PNs (baseline: 10.22 ± 1.366 Hz; ACET: 16.10 ± 3.003 Hz, paired t-test, t=2.661, df=7, *p=0.0324; not shown), due to enhanced activity of somatostatin expressing interneurons upon release of PV neuron-mediated inhibition (Englund et al., 2021). In the presence of ACET, the light-induced increase in sIPSC frequency, reflecting asynchronous GABA release from PV interneurons, was significantly smaller as compared to the control conditions (Figure 3B, C). Together, these data indicate that presynaptic KARs in PV interneurons are endogenously active and facilitate both action potential-dependent and asynchronous GABA release.

### Chronic stress associates with loss of GluK1 expression and function in LA PV interneurons

Our data so far indicate that GluK1 KARs, facilitating GABA release from PV interneurons, maintain tonic GABA_B_-mediated inhibition of the LA PNs. In order to understand whether this mechanism is regulated by chronic stress, we went on to investigate whether CRS affects GluK1 expression in PV interneurons in the LA. Since the antibodies against GluK1 subunit of KARs have low specificity in brain sections, we performed triple in situ hybridization (ISH) using fluorescent probes against *Grik1* (GluK1), *Pvalb* (parvalbumin) and *Gad1* (GAD67, marker of the GABAergic neurons).

In control sections, *Grik1* ISH signal was detected in most *Gad1* positive GABAergic interneurons (*Grik1+Gad1+/Gad1+,* 94 ± 1 %). *Pvalb* expressing neurons represented a subpopulation (23 ± 2 %) of the *Gad1* expressing neurons, which typically co-expressed *Grik1* (*Grik1+Pvalb+/Pvalb+,* 96 ± 2 %) (Figure 4A).

**Figure 4.**
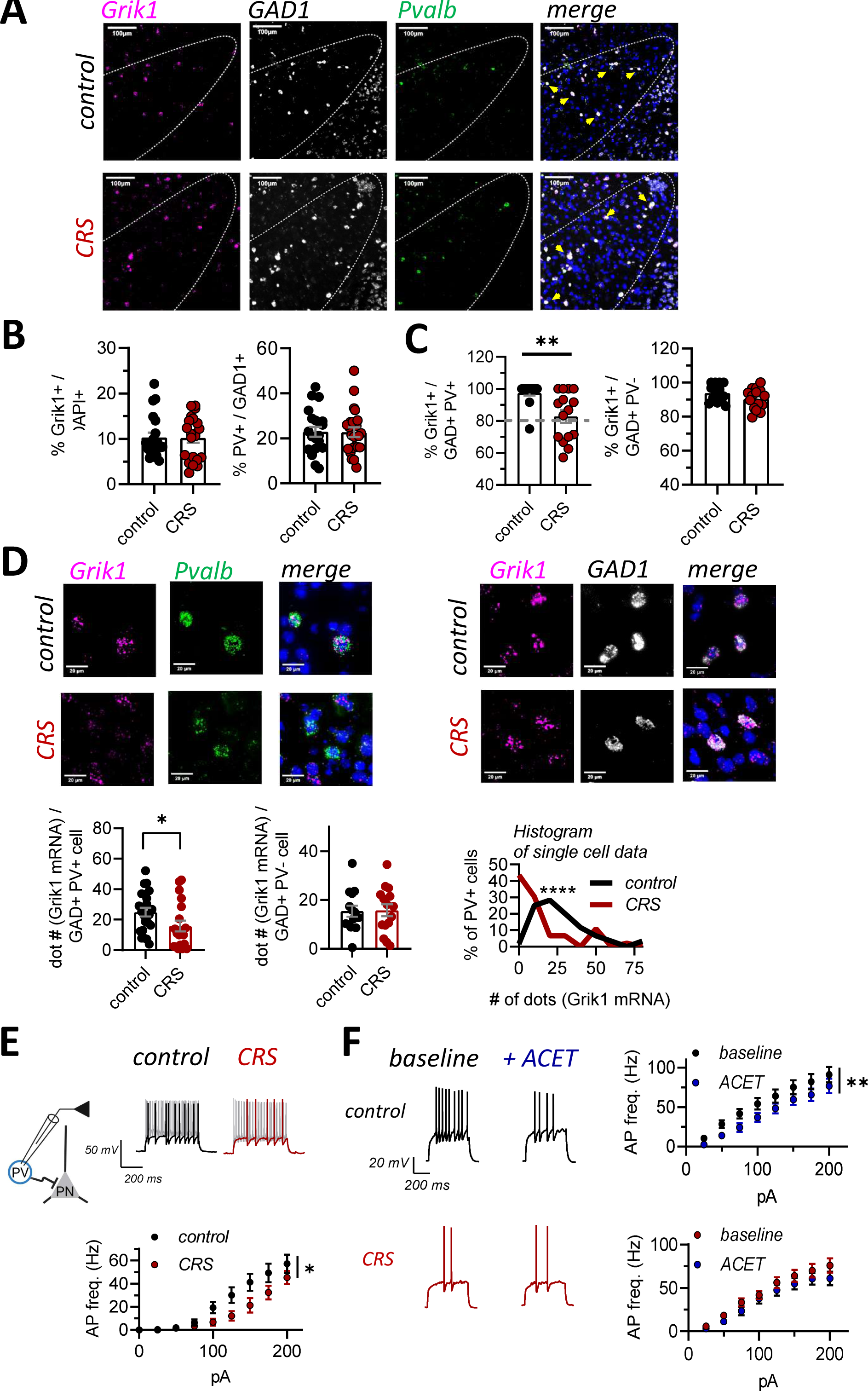
*Grik1* expression and function in PV+ interneurons is downregulated after CRS (A) Representative images illustrating triple in situ hybridization staining for *Grik1* (magenta), *Gad1* (white), and *Pvalb* (green) in the LA in control and CRS-treated mice. The image with merged channels also shows DAPI staining (blue). Yellow arrows in the merged image point to cells co-expressing *Grik1, Gad1* and *Pvalb*. (B) Bar charts summarizing the density of cells expressing *Grik1* and *Pvalb* in the LA in control and CRS- treated mice. The *Grik1* expression level is expressed as a percentage of DAPI stained nuclei, and *Pvalb* as a percentage of all *Gad1* positive GABAergic cells (control, n=20 sections (3 mice), CRS, n=21 sections (3 mice), *Grik1:* t-test, t=0.04723, df=39, p= 0.9626; *Pvalb*: t=0.05947, df=42 p=0.9529). The values represent mean ± SEM. (C) Bar charts summarizing the percentage of PV neurons (*Pvalb*+*Gad1*+) and other subtypes of GABAergic interneurons (*Pvalb-Gad1*+) co-expressing *Grik1* mRNA in LA of control and CRS-exposed mice (*Pvalb*+: control, n=14 section (3 mice), CRS, n=15 (3 mice), Mann-Whitney U= 38.50, **p=0.001; *Pvalb-Gad1*+: control, n=13 section (3 mice), CRS, n=15 (3 mice), GAD: t=1.594, df=26, p= 0.1230). The values represent mean ± SEM. (D) High-magnification images illustrating the expression of *Grik1* mRNA (magenta) in individual *Pvalb* positive LA neurons (left), and in *Pvalb* negative, *Gad1* expressing neurons (*Pvalb*, green, *Gad1*, white). The image with merged channels also shows DAPI staining (blue). Pooled data summarizing the average intensity of *Grik1* mRNA staining (# dots) per cell, in PV neurons (*Pvalb*+) and in other subtypes (*Pvalb-Gad1*+) of GABAergic interneurons in LA of control and CRS animals (*Pvalb*+: control, n=22 sections from 3 animals, CRS, n=19 sections from 3 animals, Mann-Whitney test, U=128, *p=0.034; *Pvalb-Gad1*+: control, n=14 sections from 3 animals, CRS, n=15 sections from 3 animals, Mann-Whitney test, U=100.5, p= 0.8554). Data are presented as median and quartile. Histogram demonstrating the distribution of *Grik1* mRNA staining intensity in individual PV neurons in control and CRS mice (control, n=60 cells, CRS, n=46 cells, 3 animals in both groups, Kolmogorov-Smirnov test, D= 0.4841, ****p<0.0001). (E) Current clamp recordings from PV interneurons in acute slices from control and CRS-exposed PV- TdTomato mice. Example traces illustrate the response of the PV neurons to depolarizing currents steps (50 pA and 200 pA), for control and CRS groups. Pooled data on the action potential frequencies in response to depolarizing current steps (control, n=17 (7 mice), CRS, n=18 (6 mice), RM ANOVA F_(1, 32)_ = 4.370, *p=0.0446) (F) Example traces illustrate the response of the PV neurons before and after ACET application (50 pA current step), in control and CRS-exposed PV-TdTomato mice. Pooled data on the action potential frequencies in response to depolarizing current steps (control: n=13 (6 mice), F_(1, 12)_ = 11.36, **p=0.0056; CRS: n=11 (6 mice), RM ANOVA F_(1, 10)_ = 4.112, p=0.0701) All the data presented as mean ± SEM. Data on resting membrane potential (Vm) and properties of the action potentials (AP) (rheobase, threshold and half-width) for LA PV interneurons in control and CRS- exposed mice is shown in Supplementary data 2.

The density of *Grik1* expressing neurons in the LA was not significantly different between CRS-exposed and control mice (Figure 4B). Also, the relative density of the *Pvalb* expressing GABAergic neurons in the LA was not affected by CRS treatment (Figure 4B). However, the percentage of *Pvalb* neurons co-expressing *Grik1* was significantly lower in CRS group as compared to the controls, while no difference in *Grik1* expression was detected in other subtypes of GABAergic neurons, expressing *Gad1* but not *Pvalb* (Figure 4C).

For further insight into the effect of chronic stress on *Grik1* expression levels, the intensity of *Grik1* ISH signal in individual cells was quantified by counting the number of dots representing *Grik1* mRNA staining. The mean intensity of *Grik1* ISH staining in *Pvalb* positive cells was substantially lower in the CRS group as compared to controls (Figure 4D). Again, this difference was not observed in other subtypes of GABAergic neurons, expressing *Gad1* but not *Pvalb* (Figure 4D). These data indicate that chronic stress results in downregulation of *Grik1* expression selectively in LA PV interneurons.

To test whether the observed downregulation of *Grik1* expression was sufficient to perturb GluK1 KAR function, we did current clamp experiments from LA PV interneurons in acute slices from control and CRS- exposed PV-TdTom mice. As shown previously (Englund et al., 2021), KAR antagonism by ACET resulted in significant attenuation of the PV interneuron firing rate in response to depolarizing current steps in slices from control mice (Figure 4E, F). In CRS-exposed mice, the basal excitability of the PV interneurons was significantly lower as compared to the controls, and ACET application had no significant effect on the firing frequency (Figure 4E, F). These data indicate that CRS results in loss of functional KARs and reduced excitability in LA PV interneurons.

### Chronic stress and loss of the GluK1-GABA_B_-dependent tonic inhibition regulates LA output to CeL in a cell- type specific manner

LA principal neurons have direct and indirect projections to the centrolateral amygdala (CeL), containing functionally distinct subpopulations of GABAergic neurons that can be identified based on expression of specific marker proteins (SOM, PKCδ and CRF; Moscarello and Penzo 2022). Plastic changes in the intra- amygdaloid connectivity can define activation of distinct output pathways underlying fear-related behaviors (Li et al., 2013; Hartley et al., 2019) yet it remains unclear how the circuitry is modulated by chronic stress. To understand how LA hyperexcitability in response to loss of the GluK1-GABA_B_-mediated tonic inhibition and chronic stress affects LA output to CeL, we focused on the connections to the CeL PKCδ expressing (PKCδ+) neurons implicated in the regulation of anxiety-like behaviors (Cai et a., 2014; Botta et al., 2015).

Recordings of spontaneous glutamatergic activity (sEPSCs) in slices from PKCδ-TdTom mice indicated that the basal sEPSC frequency in PKCδ+ neurons was higher in CRS-exposed mice as compared to controls, while in the surrounding PKCδ-CeL neurons, the sEPSC frequency was not significantly affected by the CRS treatment (Figure 5A). We also analyzed the frequency of sIPSCs from the same recordings (Supplementary Figure 3A), but found no differences between the groups. CeL neurons receive glutamatergic inputs from several brain areas. To test whether the observed increase in the sEPSC frequency in the PKCδ+ neurons depended on BLA hyperexcitability after chronic stress, we used chemogenetic tools to selectively inhibit activity of BLA PN neurons in control and CRS-exposed mice (Figure 5B). CNO application in acute slices from mice expressing the inhibitory DREADD receptor hM4Di in the BLA PNs resulted in 49 ± 8% reduction in the sEPSC frequency in PKCδ+ neurons, indicating that BLA neurons significantly contribute to the excitatory drive to PKCδ+ neurons. When the same experiment was repeated in CRS-treated mice, the % inhibition of the sEPSC frequency in response to CNO application was significantly higher than in controls (71± 5%, t-test vs control group, p=0.046). Importantly, in the presence of CNO, sEPSC frequency in PKCδ+ neurons was not different between the control and CRS groups (Figure 5C). These data indicate that chronic stress-associated BLA hyperexcitability shifts the balance of excitatory drive towards PKCδ+ cells in the CeL.

**Figure 5.**
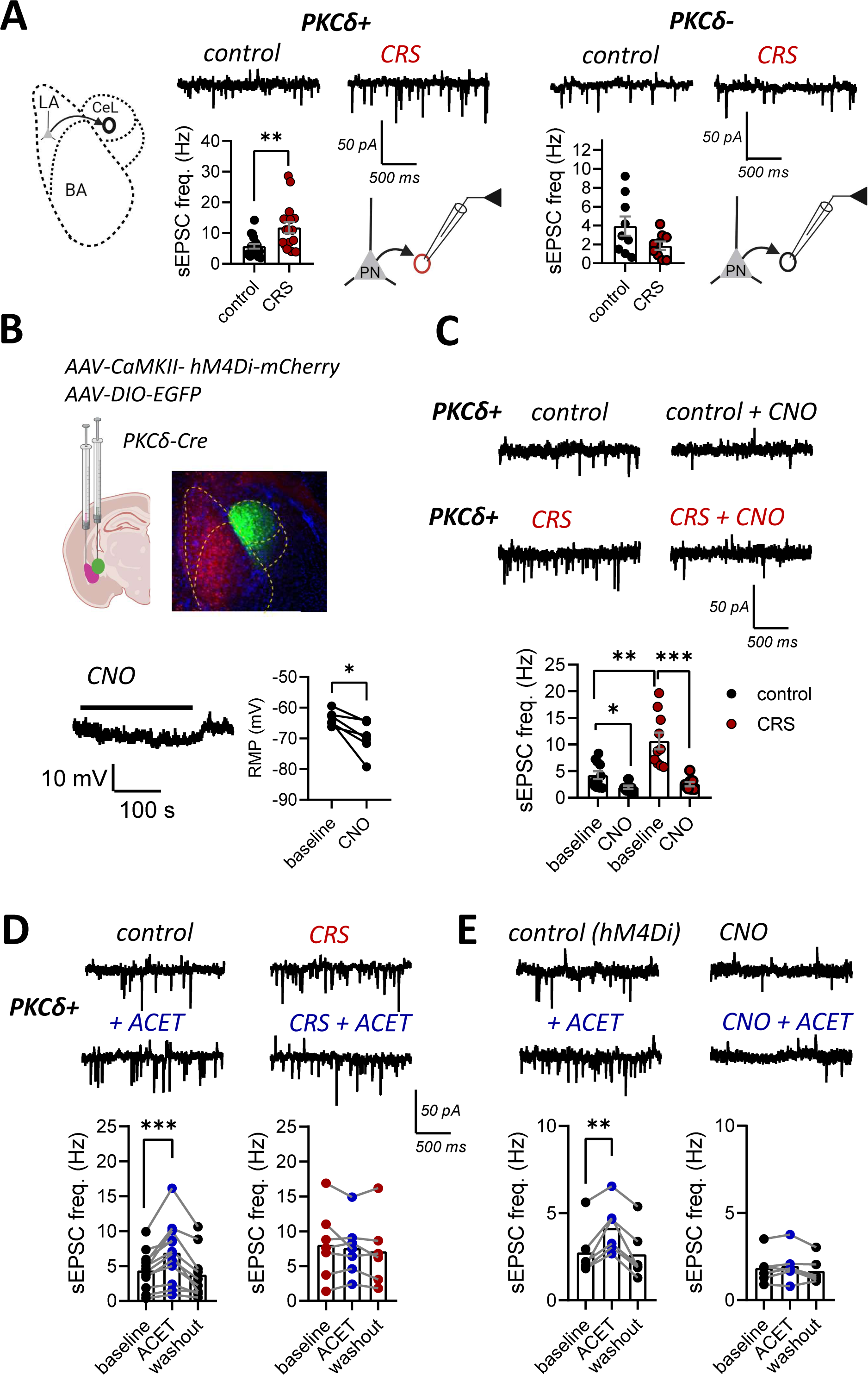
Chronic stress and loss of the GluK1-GABA_B_-dependent tonic inhibition regulates LA output to CeL (A) Voltage clamp recordings of sEPSCs from PKCδ+ and PKCδ- neurons in CeL, in acute slices from control and CRS-exposed PKCδ-TdTom mice. Pooled data on the sEPSC frequency in the two cell types (PKCδ+: control, n=19 (7 mice), CRS, n=16 (5 mice), t-test, t=3.302, df=33, **p=0.0023; PKCδ-: control, n=9 (4 mice), CRS, n=9 (5 mice), t-test, t=1.848, df=6, p=0.1140). (B) The experimental approach for chemogenetic inhibition of the BLA PNs in PKCδ-Cre mice. AAV viral vectors encoding for inhibitory DREADD receptor hM4Di under the CaMKII promoter were injected into the BLA to target PNs. PKCδ+ neurons in the CeL were visualized by injection of AAV viral vectors encoding Cre-dependent EGFP. sEPSCs were recorded from EGFP positive PKCδ neurons in the CeL. The effect of CNO (10 μM) on resting membrane potential (RMP) of hM4Di expressing BLA PNs (n=6 (3 mice), paired t-test, t=3.468, df=5, *p=0.0179). (C) Examples of sEPSC recordings from PKCδ+ neurons in slices from control and CRS-exposed mice, expressing the inhibitory DREADD receptor hM4Di in the BLA PNs, at control conditions and in the presence of CNO (10µM). Pooled data on the effect of CNO on sEPSC frequency (control: baseline, n=11, CNO, n=10 (4 mice), paired t-test, t=2.746, df=19, *p=0.0128; CRS: baseline, n=10, CNO, n=8 (4 mice), paired t-test, t=4.493, df=16, ***p=0.0004). Comparisons between control vs. CRS: unpaired t-test, t=3.836, df=19, **p=0.0011; control + CNO vs. CRS + CNO: unpaired t-test, t=1.204, df=16, p=0.2463. (D) Example traces and pooled data on sEPSC recordings from PKCδ+ neurons before and after application of ACET (200nM) in control and CRS-exposed animals (control: n=12 (5 mice), paired t- test, t=4.452, df=11, ***p=0.001; CRS: n=7 (5 mice), paired t-test, t=0.7174, df=6, p=0.50) (E) Examples of sEPSC recordings from PKCδ+ neurons before and after application of ACET (200nM) in mice expressing hM4Di in the BLA, in absence or continuous presence of CNO. Pooled data on the effect of ACET on sEPSC frequency (hM4Di: n=6 (4 mice), paired t-test, t=4.881, df=5, **p=0.0045; hM4Di + CNO: n=6 (4 mice), paired t-test, t=1.447, df=5, p=0.2074). All the data presented as mean ± SEM

Finally, we investigated whether the loss of GluK1 KARs, regulating the excitability of the LA PNs via tonic inhibition contributed to the increase in the sEPSC frequency in PKCδ+ neurons after chronic stress. To this end, we tested the effect of GluK1 antagonist ACET on sEPSCs in PKCδ+ neurons in control and CRS-treated mice, and during DREADD-mediated inhibition of BLA neurons. Application of ACET resulted in a significant increase in sEPSC frequency in PKCδ+ neurons in the control group, but had no effect when the same experiment was performed after CRS treatment (Figure 5D). Furthermore, ACET application during chemogenetic inhibition of BLA PN’s had no effect on sEPSC frequency in PKCδ+ neurons (Figure 5E), suggesting that the ACET-induced increase in glutamatergic drive to PKCδ+ neurons was due to altered excitability of BLA PNs. Since KARs are implicated in regulation of glutamate release in the BLA-CeL synapses (Arora et al., 2018), particularly in young animals (Ryazantseva et al., 2020), we also tested whether pharmacological manipulation of GluK1 KARs affected glutamate release probability, by recording the paired- pulse facilitation ratio (PPR) of EPSCs in PKCδ+ neurons, in response to stimulation of the LA. However, in contrast to the neonatal animals (Ryazantseva et al., 2020), application of GluK1 agonist ATPA or ACET had no effect on PPR in the adult mice (Supplementary Figure 3B). Together, these data indicate that stress-induced loss of the GluK1–GABA_B_-mediated tonic inhibition of the LA PNs facilitates glutamatergic drive selectively to the PKCδ+ neurons in the CeL.

### Mice lacking *Grik1* expression in PV interneurons are resistant to CRS-induced alterations in amygdala excitability and behavior

In order to understand the significance of the GluK1-GABA_B_-mediated regulation in stress-induced amygdala hyperexcitability and anxiety-like behaviors, we exposed the mice lacking GluK1 expression selectively in PV interneurons (*PV-Cre*::*Grik1^fl^*^/*fl*^) together with littermate controls (*Grik1^fl^*^/*fl*^) to chronic restraint stress. Both strains showed a significant loss of weight during the stress protocol, indicative of a physiological response to the stress (Figure 6A).

**Figure 6.**
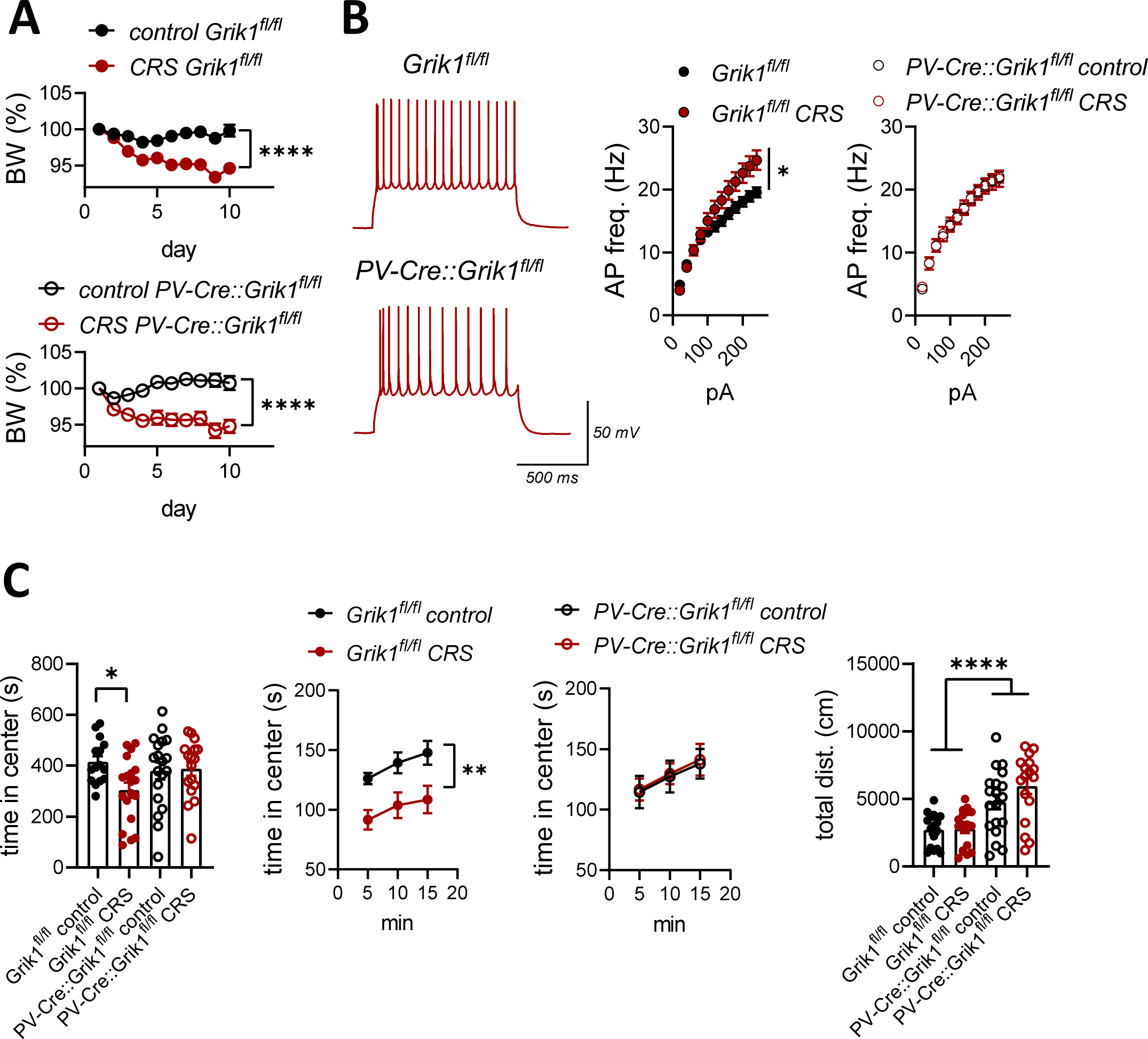
Mice lacking Grik1 expression in PV+ interneurons are resistant to stress-induced alterations in amygdala excitability and behavior (A) Body weight changes during restraint stress protocol (Grik1^fl/fl^: stress, n=19; control, n=17; RM ANOVA, F _(1, 34)_ = 21.32, ****P <0.0001; PV-Cre::Grik1^fl/fl^: stress, n=17, control, n=19; RM ANOVA; F _(1, 34)_ = 23.89, ****P <0.0001) (B) Action potential frequencies in response to depolarizing current steps recorded from brain slices of control and CRS-treated, Grik1^fl/fl^ or PV-Cre::Grik1^fl/fl^ mice (Grik1^fl/fl^: control, n=25 (7 mice), CRS, n=17 (6 mice), RM ANOVA, F _(1, 40)_ = 4.775, *p=0.035; PV-Cre::Grik1^fl/fl^: control, n=19 (6 mice), CRS, n=14 (4 mice), RM ANOVA, F _(1, 31)_ = 0.0001180, p=0.99) (C) Results of open field (OF) test. The graphs show the total time spent in the center area of the open field (OF) and the center time in 5 min bins, for control and CRS-exposed mice for the two genotypes (Grik1^fl/fl^ and PV-Cre::Grik1^fl/fl^). Grik1^fl/fl^: control, n=16, stress, n=19; PV-Cre::Grik1^fl/fl^: control, n=19, stress, n=17. Total time in the center field: Two-way ANOVA: treatment F_(1, 67)_=3.086, p=0.0835, genotype F_(1, 67)_=0.7517, p=0.389, interaction F_(1, 67)_= 4.133, p=0.046. * p=0.019, Holm Sidak. Time in center zone in 5 min bins: RM ANOVA, Grik1^fl/fl^: F_(1, 34)_=8.564, **p=0.0062; PV-Cre::Grik1^fl/fl^ F _(1, 34)_=0.03320, p=0.8566. Total distance traveled: 2-way ANOVA, genotype F_(1,67)_=31.74, ****p<0.0001, treatment F_(1, 67)_=1.828, p=0.1780, interaction F_(1, 67)_=1.495, P=0.2442. All the data presented as mean ± SEM B. Representative traces and averaged data on the medium duration afterhyperpolarizing (AHP) current amplitude, in the LA PNs of control and CRS exposed animals (control, n=12 (5 mice), CRS, n=16 (3 mice), Mann-Whitney test, U=29, **p=0.0012) C. Representative traces and pooled data on sIPSC frequency and amplitude, recorded from PN in the LA of control and CRS exposed animals, using high-Cl containing electrode filling solution in the presence of antagonists for ionotropic glutamate receptors (control, n=12 (4 mice), CRS, n=15 (4 mice), frequency: t-test, t=1.381, df=25, p=0.1796; amplitude: t-test, t=1.276, df=25, p=0.2136) D. Superimposed traces of averaged sIPSCs and the distribution of their rise and decay time properties, for the same data as in C (decay: chi-squared, p = 0.738, rise time: chi-squared, p = 0.999). E. Outward currents in response to application of GABA_B_ receptor agonist SKF97541 (25 µM) in LA PNs of C57/BL6 mice (control, n=7 (3 mice), GDP-β-S, n=6 (3 mice); t-test, t=6.108, df=11, ****p<0.0001), and Grik1^fl/fl^ and PV- Cre::Grik1^fl/fl^ mice (Grik1^fl/fl^, n=12 (5 mice), PV-Cre::Grik1^fl/fl^, n=8 (3 mice); t-test, t=0.2039, df=18, p= 0.8407). The G- protein inhibitor GDP-β-S (750 μM) was added to the pipette solution. All the recordings were done in the presence of 50 μM of DAP-5, 200 μM picrotoxin, 50 μM GYKI 53655. The GABA_B_ receptor antagonist CGP55845 (5 µM) was added in the end of the experiment, and fully blocked the SKF97541 induced current. All the data are presented as mean ± S.E.M.

The excitability of LA PNs was investigated using current clamp recordings in acute brain slices from control and CRS-exposed mice, of both strains. As expected, in the *Grik1^fl^*^/fl^ mice, the firing rate of the LA PN’s in response to depolarizing current steps was significantly higher in slices from CRS-exposed mice as compared to controls (Figure 6B). In control mice, AP firing frequency during depolarizing current steps was not different between the genotypes (Figure 6B). However, in contrast to controls, CRS had no effect on the firing rate of LA PNs in the *PV-Cre*::*Grik1^fl/fl^*mice lacking GluK1 KARs in PV neurons (Figure 6B).

The anxiety-like behavior was tested using an open field test (OF). The CRS treatment in the controls (*Grik1^fl^*^/*fl*^) resulted in the anticipated increase in anxiety-like behavior, involving avoidance of the open space in the center area of the arena (Figure 6C). In contrast, the CRS-exposed *PV-Cre*::*Grik1^fl/fl^* mice behaved similarly as their littermate controls and there were no differences between the groups in the time spent in the center area of the open field arena (Figure 6C). In this cohort of mice, the locomotor activity was not affected by CRS treatment; however, we observed that the *PV-Cre*::*Grik1^fl/fl^* mice were more active as compared to their littermates, indicated by the total distance traveled in the OF arena during the test (Figure 6C).

## Discussion

Changes in GABAergic inhibition contribute to stress-induced anxiety in rodents (Liu et al., 2014; Pan et al., 2020; Qin et al., 2022; Botta et al., 2015), yet the detailed molecular and circuit mechanisms involved are not fully understood. Here we show that chronic restraint stress attenuates tonic GABA_B_ receptor-mediated inhibition of LA PNs, resulting in their hyperexcitability and perturbed output to CeL neurons controlling anxiety-like behaviors. Furthermore, we show that this mechanism is regulated by GluK1-driven GABA release from PV interneurons, identifying PV interneurons and GluK1 receptors as critical gatekeepers of the stress-sensitive circuits in the amygdala.

### Release of tonic GABA_B_ receptor-mediated inhibition in amygdala after chronic stress

Previously, anxiolytic effects of GABA_B_ agonists, such as baclofen, have been described in both humans and rats, and prominent anxiety-like behaviors have been reported in mice lacking GABA_B_ receptor subunits (Kumar et al., 2013; Felice et al., 2016). Our present data provides mechanistic explanation for these findings, and directly link endogenous tonic activity of GABA_B_ receptors to amygdala excitability.

Weakening of tonic inhibition can be mediated by downregulation of the postsynaptic / extrasynaptic GABA receptors or reduction in the levels of ambient extracellular GABA. While chronic stress alters the expression of various GABA_A_ receptor subunits (Maguire, 2014) and in particular, the extrasynaptic GABA_A_δ and α5 subunits in the amygdala (Qin et al., 2022; Botta et al., 2015), no changes in the postsynaptic response to GABA_B_ agonists has been observed in response to stress (Pan et al., 2020; Qin et al., 2022). Instead, we observed that chronic stress associates with downregulation of GluK1 subunit containing kainate receptors facilitating both action potential-dependent and asynchronous GABA release in PV interneurons. Selective genetic ablation or pharmacological inhibition of these receptors substantially reduced the tonic GABA_B_ receptor-mediated inhibition of LA PNs. These results support that the stress-induced weakening of the tonic GABA_B_ currents in LA PNs depends on changes in ambient GABA, due to loss of GluK1 KARs facilitating GABA release from PV interneurons.

PV interneurons have a well-characterized role in pacing and synchronizing activity of the principal neurons (Hu et al., 2014). Intriguingly, our data show that alterations in PV interneuron physiology can also induce sustained changes in PN excitability via GABA_B_ receptor-mediated tonic inhibition and this is critical for LA hyperexcitability after chronic stress. This finding may have broader significance for mechanistic understanding of the various neurological and psychiatric conditions involving PV interneuron dysfunction and altered circuit excitability (Hijazi et al., 2023; Juarez et al., 2022).

### GluK1 KARs in PV interneurons regulate amygdala excitability

GluK1 KARs have been previously implicated in stress-induced alterations in amygdala as well as in anxiety- like behaviors (Wu et al., 2007; Masneuf et al., 2014; Englund et al., 2021). However, since GluK1 KARs are expressed in both glutamatergic and GABAergic neurons in the amygdala and are found in different subtypes of GABAergic interneurons (Daw et al., 2010; Lourenco et al., 2010; Wyeth et al., 2017; Selvakumar et al., 2021), the cell types mainly responsible for GluK1-dependent modulation of amygdala excitability and amygdala dependent behaviors have remained unclear. Our present and previous (Englund et al., 2021; Haikonen et al., 2023) results provide compelling data showing that GluK1 KARs are physiologically activated in LA PV interneurons to regulate their excitability and GABA release, which is critical for gating the excitability of the behaviorally relevant circuits in the amygdala. Accordingly, the absence of this regulation in mice lacking *Grik1* expression in PV interneurons associated with resilience to stress-induced anxiety.

In a physiological context, the GluK1–GABA_B_ interplay may operate as a feedback system restricting excitability of the LA PNs during intense glutamatergic activity. In the immature hippocampus, GluK1 KARs are physiologically activated by ambient extracellular glutamate (Lauri et al., 2006; Segerstrale et al., 2010). Assuming similar mechanism in the amygdala, accumulation of extracellular glutamate during strong synchronous excitatory activity would result in enhanced activation of GluK1 receptors in PV interneurons, facilitating GABA release and GABA_B_-receptor mediated tonic inhibition of the principal neurons. Hence, together with previous findings (reviewed in Lauri et al., 2021; Mulle and Crepel 2021), our data support that GluK1 KARs mediate crosstalk between glutamatergic and GABAergic systems to control network excitability. In addition to regulating the overall excitability, the prominent effects of GluK1 KARs on PV interneuron function could also affect fast synchronization of the amygdala circuity (Randall et al., 2011; Ojanen et al., 2023), implicated in memory consolidation (Pare and Headley, 2023).

### Stress-induced target-specific plasticity in the LA-CeL circuitry

One of the main targets of BLA principal neurons is centrolateral amygdala (CeL), which is composed of intermingled populations of functionally distinct GABAergic neurons strongly implicated in regulation of anxiety and fear generalization (Tye et al., 2011; Fadok et al., 2018; Moscarello and Penzo, 2022). BLA inputs to CeL undergo target-specific plasticity during fear acquisition and extinction (Li et al., 2013; Terburg et al., 2018; Hartley et al., 2019), affecting the relative weights of excitatory input to different CeL neuron populations. Interestingly, while dynamic modulation of the connectivity is critical for fear memory acquisition and retrieval, persistent unbalance in the cell-type specific BLA–CeL connectivity associates with anxiety–like behaviors in genetically modified mouse models (Arora et al., 2018). In our experiments, anxiogenic chronic stress resulted in a shift in glutamatergic transmission towards CeL PKCδ+ neurons, and this effect depended on LA PN hyperexcitability and loss of the GluK1–GABA_B_-mediated tonic inhibition. Since activity of PKCδ neurons has been directly linked to anxiety-like behaviors in rodents (Cai et al., 2014; Botta et al., 2015), these data suggest that the specific increase in the excitability of the LA-CeL PKCδ circuit critically contributes to stress-induced anxiety. However, it should be noted that stress widely affects the brain and modulates various networks integrating external and internal factors, which can also contribute to the behavioral outcome.

## Materials and methods

### Animals

C57BL/6N-*Grik1^tm1c^*^(KOMP)Mbp^ (*Grik1^fl^*^/fl^; Ryazantseva et al., 2020) mouse line was crossed with a B6.129P2- *Pvalb^tm1^*^(cre)Arbr/J^ (PV-Cre) (JAX 008069) line to produce ablation of *Grik1* gene selectively in the parvalbumin interneurons (PV-Cre::*Grik1^fl^*^/fl^). PKCδ-Cre mouse line (C57Bl/6CRL-*Prkcd*-*iCre* (GENSAT 011559-UCD), kindly provided by Wulf Haubensak) was crossed with Ai14 tdTomato reporter (JAX 007914) to visualize PKCδ cells in the central amygdala. PV-Cre mice were crossed with Ai14 tdTomato reporter to visualize parvalbumin interneurons in BLA. The mice were group housed in individually ventilated cages with a 12 light/12 dark cycle (lights are off 7:00 p.m. – 7:00 a.m.), and food and water were supplied *ad libitum*. All animal experiments were performed by following the University of Helsinki Animal Welfare Guidelines and approved by the National Animal Experiment Board of Finland (license numbers: KEK-17-019, KEK22-010, ESAVI/29384/2019, and ESAVI/31984/2022). Three-month-old male C57BL/6JRccHsd mice were used for ISH. Two- to three-month-old male mice were used for behavioral testing and electrophysiology.

### Chronic restraint stress (CRS)

Mice were randomly assigned into two groups: the CRS group and the control group. Littermates were placed equally in both groups. The mice in both groups were weighed daily at the same time before starting the CRS procedure. The mice belonging to the CRS group were restrained in well-ventilated 50 ml Falcon tubes at the same time each day (8:30 a.m. – 9:30 a.m.) for 10 consecutive days. During the restraining period, the control mice remained in their home cages in the same room where the restraint stress procedure took place. Food and water were supplied *ad libitum* in home cages.

### Behavioral testing

The open field (OF) test was done on the 11th day of the experiment (after the 10 days CRS protocol). The test was done in a room with dimmed lights, in four 50 cm × 50 cm arenas with 32 cm high walls, with one mouse in each arena per trial. At the beginning of each trial, one mouse was placed in the corner of each arena and observed for 15 minutes using EthoVision XT15 (Noldus, Wageningen, Netherlands) video-tracking software. At the end of each trial, the arena was cleaned with 70 % EtOH and wiped dry to remove any scent traces.

### Viral injections

AAV viral vectors encoding for inhibitory DREADD (AAV8, pAAV-CaMKIIa-hM4D(Gi)-mCherry; Addgene #50477), floxed EGFP (AAV8; pAAV-hSyn-DIO-EGFP; Addgene #50457) were bilaterally injected to the BLA of adult (>P55) heterozygote PKCδ-Cre mice. AAV viral vector encoding the Cre-activated expression of ChR2 (AAV1; pAAV-EF1a-double floxed-hChR2(H134R)-mCherry-WPRE-HGHpA; Addgene #20297) was injected bilaterally in the BLA of adult heterozygote PV-Cre mice. Injections were done under anesthesia in a stereotaxic frame as described in Ryazantseva et al., 2020, using the following coordinates for BLA (from bregma): (1) AP −1,8 ML 3.4-3.5 DV 4.1–4.3 (2) AP −2.3 ML 3.4-3.5 DV 4.1–4.3; and for CeA: AP 1.8 ML 2.9 DV 4.4.

### Electrophysiology

#### Acute slices

Acute coronal sections were prepared as previously (Englund et al., 2021). Briefly, the brain was removed and immediately placed in carbonated (95% O_2_/ 5% CO_2_) ice-cold N-Methyl-D-glucamine (NMDG) based protective cutting solution (pH 7.3–7.4) containing (in mM): 92 NMDG, 2.5 KCl, 1.25 NaH_2_PO_4_, 30 NaHCO_3_, 20 HEPES, 25 glucose, 2 thiourea, 5 Na-ascorbate, 3 Na-pyruvate, 0.5 CaCl_2_ and 10 MgSO_4_ (Ting et al., 2014). The vibratome (Leica VT 1200S) was used to obtain 300-µm-thick brain slices. Slices containing the amygdala were placed into a slice holder and incubated for 8–10 min in 34 °C in the NMDG–based solution. Slices were then transferred into a separate slice holder at room temperature with a solution containing (in mM): 92 NaCl, 2.5 KCl, 1.25 NaH_2_PO_4_, 30 NaHCO_3_, 20 HEPES, 25 glucose, 2 thiourea, 5 Na-ascorbate, 3 Na-pyruvate, 2 CaCl_2_ and 2 MgSO_4_ (saturated with 95% O_2_/5% CO_2_).

#### Whole-cell recordings

After 1–4 h of recovery, the slices were placed in a submerged heated (30-32 °C) recording chamber and continuously perfused with standard ACSF, containing (in mM): 124 NaCl, 3 KCl, 1.25 NaH_2_PO_4_, 26 NaHCO_3_, 15 glucose, 1 MgSO_4_ and 2 CaCl_2_, at the speed of 1.5–2 ml/min. Whole-cell patch-clamp recordings were done from amygdala neurons under visual guidance using glass capillary microelectrodes with resistance of 3–5.5 MΩ. Multiclamp 700B amplifier (Molecular Devices), Digidata 1322 (Molecular Devices) or NI USB-6341 A/D board (National Instruments), and WinLTP version 2.20 (Anderson et al., 2007) or pClamp 11.0 software were used for data collection, with low pass filter (10 kHz) and a sampling rate of 20 kHz. In voltage-clamp recordings, uncompensated series resistance (Rs) was monitored by measuring the peak amplitude of the fast whole-cell capacitance current in response to a 5 mV step. Only experiments where Rs <30 MΩ, and with <20% change in Rs during the experiment, were included in the analysis.

*Whole-cell current clamp recordings of membrane excitability* in principal neurons and PV interneurons in LA were performed using a filling solution containing (in mM): 135 K-gluconate, 10 HEPES, 5 EGTA, 2 KCl, 2 Ca(OH)_2_, 4 Mg-ATP, 0.5 Na-GTP, (280 mOsm, pH 7.2). The resting membrane potential was sampled and then adjusted to -70 mV. Depolarizing current steps with 600 ms (PV interneurons) or 1000 ms duration (principal neurons) were applied to induce action potential firing. The amplitude of the injected current was increased with 10-20 pA increments. Since the principal neuron firing rate is not stable under whole-cell recording conditions, the excitability data was always collected immediately after whole-cell access, under control conditions or in the presence and after > 10 min preincubation of a relevant antagonist (200 nM ACET, 5 µM CGP55845). The effect of GluK1 antagonism on PV interneuron excitability was addressed by using fast application of 200 nM ACET in the continuous presence of 50 µM GYKI 53655, 5 µM L-689560 and 100 µM picrotoxin, to block AMPA, NMDA and GABA_A_ receptors.

*Drug-induced GABA_A_- and GABA_B_-mediated currents* were recorded from LA PNs in the presence of antagonists for NMDA and AMPA receptors (50 µM AP-5 and 50 µM GYKI 53655, respectively). GluK1 antagonist ACET (200nM) was included in some of the experiments. For recordings of GABA_B_ receptor- mediated currents, the intracellular solution contained (in mM): 140 K-gluconate, 10 HEPES, 1 EGTA, 2 KCl, 2 NaCl, 4 Mg-ATP, 0.5 Na-GTP, (280 mOsm, pH 7.2), 100 µM picrotoxin was included in the ACSF to block GABA_A_ receptors, and the holding potential of the neurons was -50 mV. For recordings of GABA_A_-mediated currents, the intracellular solution contained (in mM): 130 CsCl, 10 HEPES, 0.5 EGTA, 8 NaCl, 4 Mg-ATP, 0.3 Na-GTP, 5 QX314 (280 mOsm, pH 7.2), 5 µM CGP 55845 hydrochloride was included in the ACSF to block GABA_B_ receptors, and the holding potential of the neurons was -90 mV.

A valve control system (VC-6-PINCH, Warner Instruments) with a manifold adjusted to the tube was used for fast drug application. ACSF including the above antagonists was applied directly to the LA to record a baseline before switching to solution supplemented with the relevant drug (25 µM CGP55845 hydrochloride, 25 µM SKF97541 or 25 µM(-)-Bicuculline methochloride).

*mAHP currents.* Medium duration afterhyperpolarizing (mAHP) currents were recorded from LA PNs under voltage clamp , at holding potential -50 mV, using filling solution containing (in mM): 135 K-gluconate, 10 HEPES, 5 EGTA, 2 KCl, 2 Ca(OH)_2_, 4 Mg-ATP, 0.5 Na-GTP, (280 mOsm, pH 7.2). mAHP currents were induced by 50 ms depolarization step from -50 mV to 0 mV.

#### Spontaneous synaptic currents

sEPSCs and sIPSCs in PKCδ neurons and LA principal neurons were recorded under whole-cell voltage-clamp with a filling solution containing (in mM): 135 K-gluconate, 10 HEPES, 5 EGTA, 2 KCl, 2 Ca(OH)_2_, 4 Mg-ATP, 0.5 Na-GTP, (280 mOsm, pH 7.2) at holding potential of -50 mV. ACET (200 nM) and CNO (10 µM) were added to the ACSF. For more detailed analysis of sIPSCs in LA principal neurons and for light-induced IPSCs, recordings were done using high-chloride intracellular solution containing (in mM): 130 CsCl, 10 HEPES, 0.5 EGTA, 8 NaCl, 4 Mg-ATP, 0.3 Na-GTP, 5 QX314 (280 mOsm, pH 7.2), at -80 mV holding potential.

*Evoked EPSCs* were recorded in from PKCδ neurons in CeA at a holding potential of −70 mV, using electrodes filled with the following solution (in mM): 130 CsMeSO_4_, 10 HEPES, 0.5 EGTA, 8 NaCl, 4 Mg-ATP, 0.3 Na-GTP, 5 QX314 (280 mOsm, pH 7.2). Picrotoxin (100 µM) and D-AP5 (50 µM) were included in the perfusion solution to antagonize fast GABA_A_ and NMDAR-mediated components of synaptic transmission, respectively. EPSCs were evoked with bipolar stimulation electrodes (nickel-chromium wire), placed at LA to mimic the activation of glutamatergic LA afferents. Paired-pulse responses (PPR) were evoked with the inter-pulse interval of 50 ms.

*Optical stimulation* of PV interneurons was done using pE-2 LED system (CoolLED, UK). Short pulses (0.2- 0.5 ms, 50 ms interval) of 470 nm blue light were used for paired-pulse stimulation . Asynchronous release was triggered with 750 ms long light pulses. LED intensity was kept constant throughout the recording.

#### Data analysis

WinLTP software (Anderson and Collingridge, 2007) was used to calculate the peak amplitude of the evoked synaptic responses. For analysis of paired-pulse ratio (PPR), 7–10 responses were averaged in each experimental condition. PPR was calculated as the amplitude ratio of response 2/response 1. The frequency and amplitude of spontaneous synaptic events were analyzed using miniAnalysis program 6.0.3. sIPSCs and sEPSCs were identified in the analysis as outward or inward currents (for IPSCs depending on experimental conditions) with typical kinetics, respectively, that were at least 3 times the amplitude of the baseline level of noise. For the pooled data, averages for baseline, drug application and washout were calculated over a 10-minute period. Action potential frequencies were analyzed using the threshold search algorithm in Clampfit software. AP half-width and amplitude of the fast hyperpolarizing potential are analyzed from the 3^rd^ spike in the train using Clampfit software.

### In situ hybridization

Triple-ISH was done using the RNAscope® Fluorescent Multiplex kit (Advanced Cell Diagnostics) on fresh frozen brain sections from control and CRS-exposed mice. The brain was removed under anesthesia, flash frozen in dry ice and cut onto 10-µm-thick coronal sections with a cryostat (Leica CM3050). The sections dried in the cryostat at -20 °C for 1 h, after which they were stored at -80 °C until the next day. Before hybridization, the sections were fixed in cold 4 % PFA for 30 min, dehydrated in ethanol series, and treated with protease for 30 min at room temperature (RT). The hybridization reaction was performed according to the manufacturers instructions, using the following probes: Mm-Pvalb-C1, Mm-GAD1-C2, and Mm-Grik1-C3 (Advanced Cell Diagnostics). After hybridization, the slices were stained with DAPI. The sections containing the BLA were imaged with Zeiss Axio Imager.M2 microscope equipped with ApoTome 2 attachment, using PlanApo ×20 and ×40 objectives, CMOS camera (Hamamatsu ORCA Flash 4.0 V2), and Zen 2 software. The density of stained cells with the brain region of interest were analyzed from ×20 images using ImageJ v1.53f software. The threshold for *Grik1* expression was assigned to >2 dots co-localized with a cell marker. The signal intensity of *Grik1* expression at the single-cell level was quantified by counting the number of puncta per cell semi-automatically with QuPath software v0.3.2 (Bankhead et al., 2017) from ×40 images. All the data were obtained from at least 4 sections/animal (4 animals per group).

### Statistical analysis

All statistical analyses were done on raw (not normalized) data using Graph Pad Prism 9.0 software. Sample size was based on previous experience on similar experiments. All data were first assessed for normality and homogeneity of variance and the statistical test was chosen accordingly. Differences between two groups were analyzed using two-tailed t-test or Mann-Whitney rank sum test. Two-way ANOVA with Holm-Sidak post hoc comparison was used to compare effects of genotype and stress. For data that was not normally distributed, Mann-Whitney test was used for post-hoc pairwise comparison. Dunnett test was used for testing two or more experimental groups against a single control group. Chi-squared test was used to compare distributions of properties for synaptic events. Kolmogorov-Smirnov test was used to compare distributions of Grik1-expressing parvalbumin interneurons in control and stressed animals’ amygdala. To compare drug effects to the baseline, Student’s paired two-tailed t-test was used. The results were considered significant when p < 0.05. All the pooled data are given as mean ± S.E.M.

## Acknowledgements

We thank Janne Sulku, Vootele Voikar, the personnel in the Mouse Behavioral Phenotyping Facility and in the Laboratory Animal Center for expert technical help. M.R. thanks Svetlana Molchanova for inspiring discussions. This study was financially supported by the Academy of Finland (#330710, SEL; #330298, MR) and Sigrid Juselius Foundation (#1158, SEL). Mouse Behavioral Phenotyping Facility is supported by Biocenter Finland.

## Author contributions

S.E.L and M.R. designed the study, M.R. performed the electrophysiological experiments and imaging with the help of M.L., M.L. performed the in situ hybridization and image analysis and contributed to the CRS treatments. S.E.L. wrote the manuscript.

The authors declare no competing interests

---

**Supplementary Figure 1.**
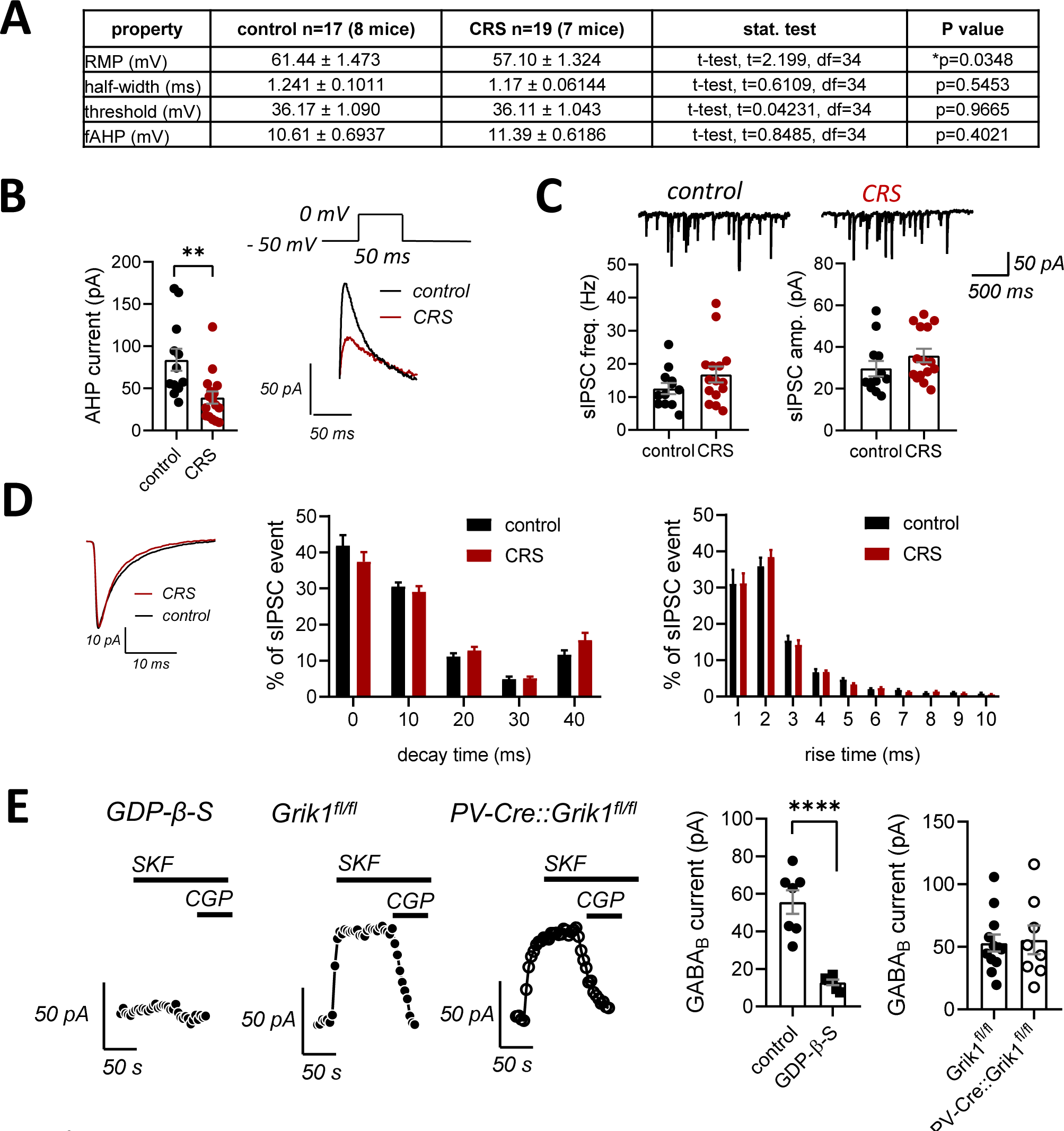
A. Membrane properties of LA principal neurons (PN) in control and CRS exposed animals. RMP – resting membrane potential, fAHP – fast afterhyperpolarizing potential.

**Supplementary Data 2.**
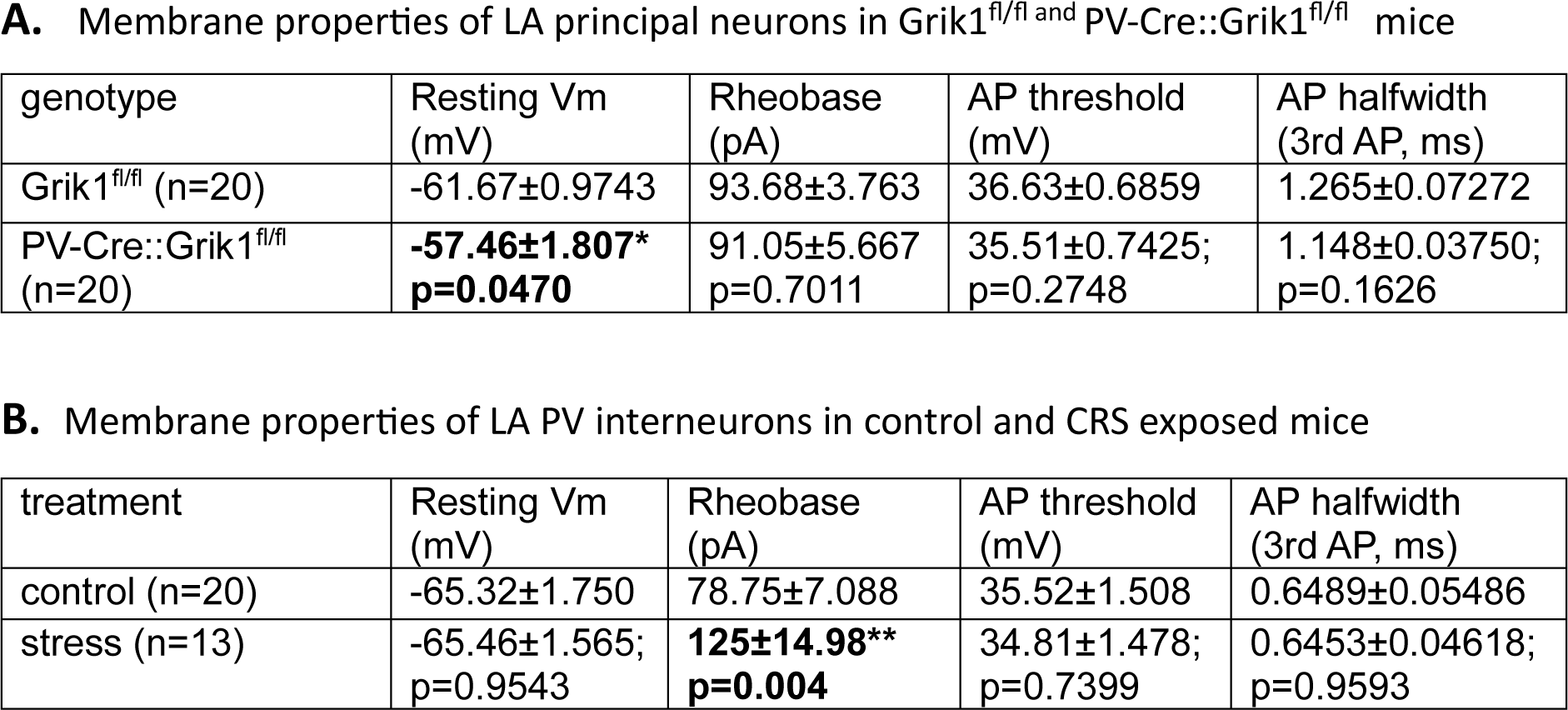
Averaged data on resting membrane potential (Vm) and properties of the action potentials (AP) (rheobase, threshold and halfwidth) for LA principal neurons in Grik1^fl/fl^ ^and^ PV-Cre::Grik1^fl/fl^ mice (A) and for LA PV interneurons in control and CRS exposed mice (B).

**Supplementary Figure 3.**
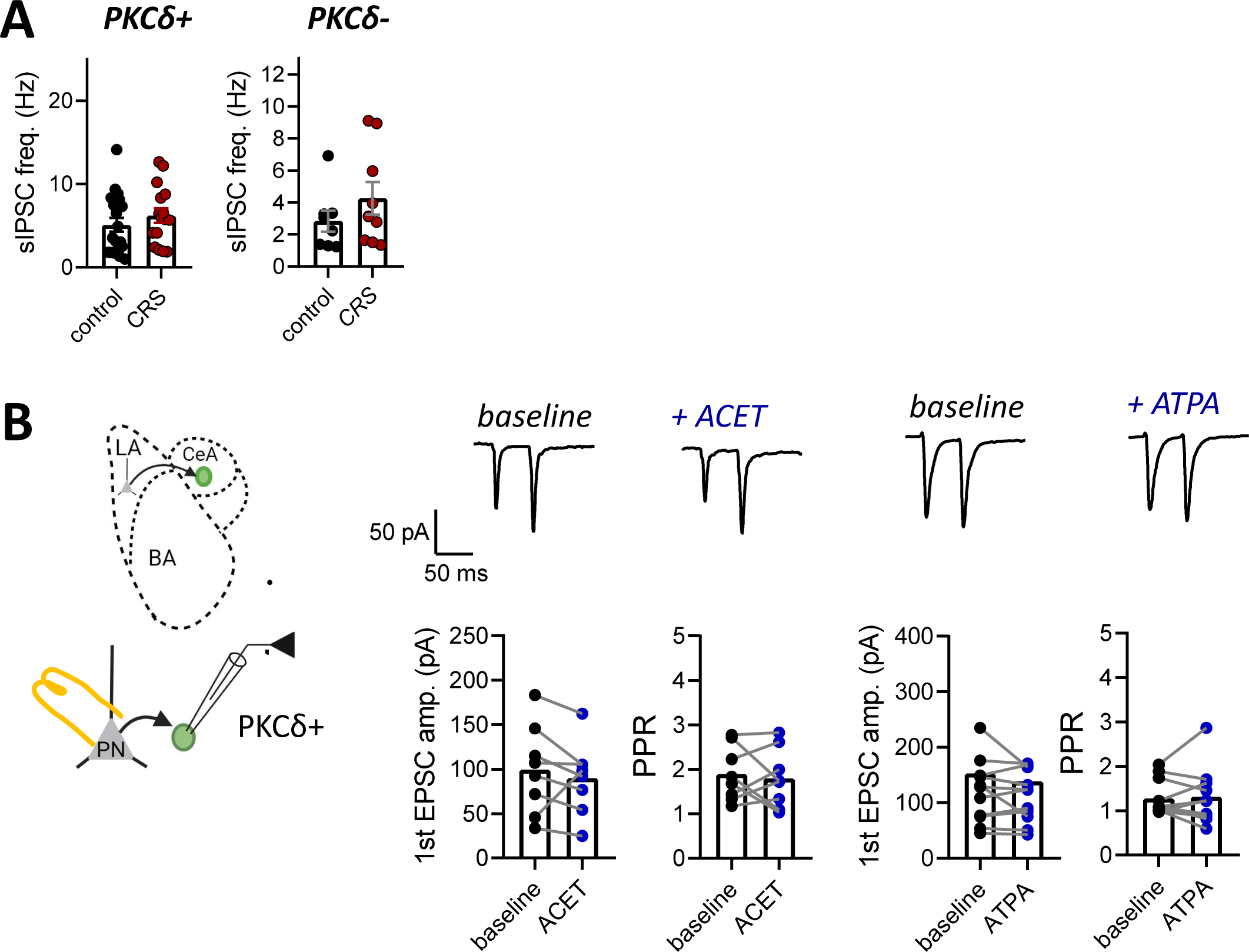
**A.** Pooled data on the sIPSC frequency , for the same recordings as shown in the Figure 5A (PKCδ+: control, n=19 (5 mice), CRS, n=16 (4 mice), t-test, t=0.9045, df=33, p=0.3723; PKCδ-: control, n=9 (4 mice), CRS, n=9 (5 mice), t-test, t=0.1988, df=16, p=0.8449). **B.** Effect of ACET (200 nM) and ATPA (1 μM) on the amplitude and paired-pulse ratio of EPSCs in CeL PKCd+ neurons, evoked by stimulation of LA (1^st^ EPSC amplitude: ACET: t=0.9624, df=7, p=0.3679; ATPA: t=1.503, df=11, p= 0.1610; PPR: ACET, n=8 (3 mice), paired t-test, t=0.4782, df=7, p= 0.6471; ATPA, n=12 (5 mice), paired t-test, t=0.3823, df=11 p=0.7095). All the data are presented as mean ± S.E.M.

